# Selective metal extraction by biologically produced siderophores during bioleaching from low-grade primary and secondary mineral resources

**DOI:** 10.1101/2021.01.15.426802

**Authors:** Adam J. Williamson, Karel Folens, Sandra Matthijs, Yensy Paz Cortez, Jeet Varia, Gijs Du Laing, Nico Boon, Tom Hennebel

**Affiliations:** Center for Microbial Ecology and Technology (CMET), Faculty of Bioscience Engineering, Ghent University, Coupure Links 653, 9000 Ghent, Belgium; SIM vzw, Technologiepark 48, 9052 Zwijnaarde, Belgium; Institut de Recherche LABIRIS, Av. E. Gryzon 1, 1070 Brussels, Belgium; Department of Green Chemistry and Technology, Faculty of Bioscience Engineering, Ghent University, Coupure Links 653, 9000 Ghent, Belgium; Umicore, Broekstraat 33, Brussels, Belgium

**Keywords:** Waste processing, Metal complexation, Pyoverdine, *Pseudomonas putida*, Resource recovery

## Abstract

Siderophores are a class of biogenic macromolecules that have high affinities for metals in the environment, thus could be exploited for alternate sustainable metal recovery technologies. Here, we assess the role of siderophores in the extraction and complexation of metals from an iron oxide-rich metallurgical processing residue and a low-grade primary Ni ore. Evaluation of the biological siderophore bioproduction by three pseudomonads, *P. fluorescens, P. azotoformans* and *P. putida* identified that *P. putida* could generate the highest siderophore yield, which was characterized as a hydroxamate and catecholate mixed-type pyoverdine PyoPpC-3B. Key physiochemical parameters involved in raw siderophore mediated metal extraction were identified using a fractional factorial design of experiments (DOE) and subsequently employed in purified PyoPpC-3B leaching experiments. Further targeted experiments with hydroxamate and catecholate functional analogues of PyoPpC-3B confirmed their marked ability to competitively or selectively leach and chelate hard metal ions, including Al(OH)_4-_, Mn^2+^ and Zn^2+^. Interestingly, complexation of Mn and Zn ions exceeded the natural affinity of pyoverdine for Fe^3+^, thus despite the low metal recoveries from the materials tested in this study, this work provides important new insights in siderophore-metal interactions.

## 1. Introduction

Siderophores are an important group of secondary metabolites produced by microorganisms and plants to facilitate the uptake of iron, which is typically insoluble in most terrestrial environments ^1–3^. Siderophore concentrations in the environment typically lie in the µM-mM range ^4^ and are intrinsically involved in weathering soil minerals ^5,6^, thus can significantly contribute to the mobility of metals in the environment. Siderophores also form complexes with a diverse range of other metals including Al, Cd, Cu, Ga, In, Pb, REE, Zr, Hf ^7,8^, essential macronutrients Mo, Mn, Co and Zn ^9^, as well as radionuclides U, Np, Th and Pu ^10–12^. Whilst the evolutionary reasoning behind this remains unclear, it could represent the sequestration of essential macronutrients, or detoxification of metals which would otherwise result in oxidative cellular stress ^9,12^.

Over 500 siderophores have been characterized to date, and can be classified by their ligand functionalities ^13^: (i) catecholates (aryl caps) of which include phenolates, (ii) hydroxamates, (iii) carboxylates or hydroxycarboxylates ^14^. It is widely documented that *Pseudomonas* sp. can produce pyoverdine-type siderophores, and less complex siderophores (‘secondary siderophores’) such as pyochelin, pseudomonine, thioquinolobactin and pyridine-2,6-bis(monothiocarboxylic acid), yet there is a paucity of studies towards the characterisation of their metal binding properties in mixed element systems. Pyoverdines have been shown to have high affinities for a range of metals including Zn, Cu and Mn (K_a_ 10^17-22^) yet with a clear preference for iron (K_a_ 10^32^) under their respective experimental conditions ^9,15,16^.

Siderophores have received much attention in recent years because of their potential application in various areas of environmental research, including medicine (e.g. anemia treatments), agriculture (plant-bacteria synergism and bio-pesticides) ^17,18^, bio-sensors, chelating agents and bio-remediation ^14^ (Table 1). Siderophores can also offer perspectives for recovering raw materials from sustainable metal reserves. The depletion of high-grade mineral resources at a reasonable accessibility (<1 km depth) has forced the mining industry to search for alternative processes that exploit low-grade mineral deposits and avoid a high energy consumption. This is further exacerbated by the predicted exhaustion of Zn, Ga, Ge, As, Rh, Ag, In, Sn, Sb, Hf, Pb, Mn and Au within 50 years, Ni, Cu, Cd, Tl, Fe and U within 100 years and platinum group metals in 150 years, based on current consumption rates ^19^. Metal recovery from alternate low-grade primary and secondary sources provides a great opportunity to meet the demand of raw materials. Zinc refining operations, for instance, have been generating large amounts of iron oxide-rich jarosite and goethite wastes, posing serious environmental, social and economic difficulties ^20^. Primary laterite ores are increasingly being investigated due to their abundance and significant quantities of important metals Co and Ni ^21^. Over the last century, bioleaching is being increasingly investigated as a more sustainable mode of hydrometallurgical metal extraction ^22^. Recent work has implicated the involvement of siderophores in the leaching of metals from fayalite slags ^23,24^ and chromite tailing ^25^ by *P. aeruginosa* and *P. putida*, however, no further siderophore characterisation or metal-siderophore interactions were assessed.

**Table 1.**
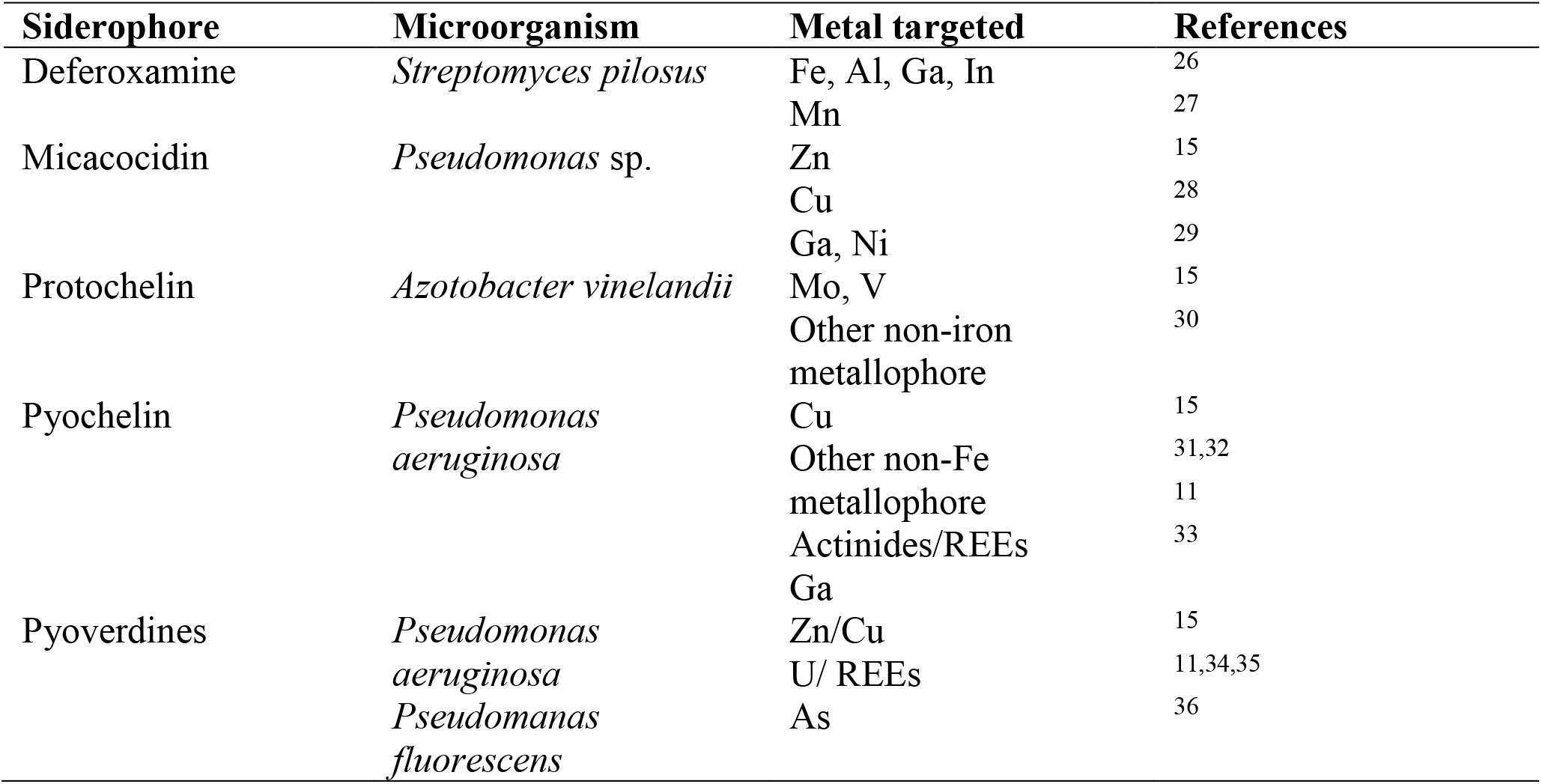
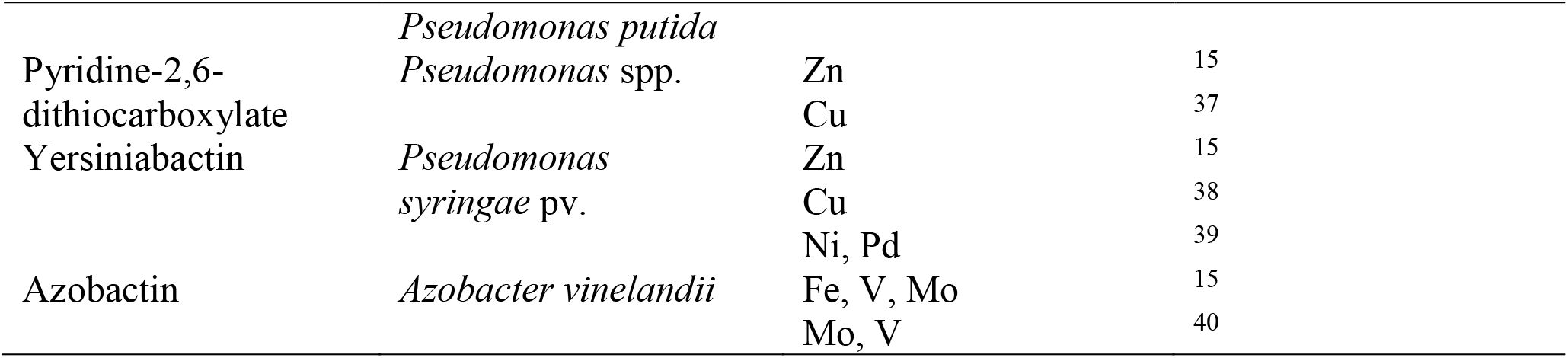
Overview of studies towards the application of siderophores towards non-Fe metals

Accordingly, this study aimed at evaluating the production of siderophores by strains of *Pseudomonas* and their potential to extract metals, including Zn, Mn and Al, from two low-grade mineral resources; the first, an iron oxide-rich residue from zinc processing, and secondly, a Ni-bearing laterite ore. Whilst improving our mechanistic understanding of siderophore-metal interactions in complex mineralogical environments, it contributes to the early development of alternate bio-metallurgical technologies for sustainable metal extraction.

## 2. Materials and Methods

### 2.1 Cultivation of strains

The bacterial strains used in this study were *Pseudomonas putida* PpF1 (LMG 24210), *Pseudomonas fluorescens* (LMG 1794) and *Pseudomonas azotoformans* (DSMZ 18862). *P. putida* and *P. fluorescens* strains were obtained from the Belgian Coordinated Collection of Micro-organisms and *P. azotoformans* was purchased from Leibniz Institute DSMZ-German Collection of Microorganisms and Cell Culture. Bacterial strains where first plated on LB agar (Carl Roth, Germany) from glycerol stocks and incubated at 28 °C for 24 h. Single colonies were further sub-cultured in 10 mL of LB broth (Carl Roth, Germany) and incubated at 28 °C for 24 h with constant shaking at 120 rpm until an optical density at 600 nm, OD_600_ (DR Lange ISIS 900 MPA photometer) was approximately 1.5. All microbial cultivation and siderophore production experiments were carried out under strictly sterile and aerobic conditions.

### 2.2 Microbial siderophore purification and characterization

To stimulate the production of siderophores, strains were first grown on LB growth media, centrifuged and washed twice with 0.9 % NaCl before transferring at a starting OD_600_ of 0.02 into a modified selective medium (MSM) previously applied for siderophore production (6 g L^-1^ K_2_HPO_4_, 3 g L^-1^ KH_2_PO_4_, 5 g L^-1^ (NH_4_)_2_SO_4_, 0.2 g L^-1^ MgSO_4_, 4 g L^-1^ Na-succinate and 4 g L^-1^ casamino acids) ^41^. The pH was set to 7 prior to autoclaving and to avoid precipitation, the casamino acids solution was filter-sterilised and added after autoclaving the medium. Strictly Fe-free conditions were established and maintained by pre-washing all glassware in 2 (v/v)% HCl, to prepare the medium in order to maximize the siderophore production ^42^. Before inoculation from LB sub-cultures to MSM, cells where centrifuged at 5000 rpm for 2 min and washed twice with MSM. The growth of each strain was evaluated in triplicate serum flasks by measuring the OD_600_ for a period of 5 days. The siderophore concentration was approximated with a high throughput chrome azurol sulfonate (CAS) assay and measuring the UV-VIS absorbance at 620 nm ^43^. The resulting pyoverdine siderophore was purified from 4 L of a 72 h old culture by previously described methods ^44^. Briefly, the filtered culture supernatant was loaded onto a C-18 column that was activated with methanol and washed with distilled water. Elution was performed with acetonitrile/ H_2_O (70/30 %). Preparative-scale purification of the pyoverdine was performed using a Prep 150 LC system (Waters). A SunFire Prep C18 column (C-18, 19 x 250 mm, 5 µm particle size) was used with a flow rate of 20 mL min^-1^ and a gradient from H_2_O/CH_3_CN 9:1 containing 0.1 % CF_3_COOH to H_2_O/CH_3_CN 6:4 containing 0.1 % CF_3_COOH in 20 min. CH_3_CN was evaporated from the extract in vacuo and the sample was lyophilized. LC/MS analyses were performed to identify the pyoverdine on a Kontron 325 system, coupled to the mass spectrometer and equipped with a UV detector (model 322), an automatic injector (model 465) and LC-6A pumps. The column used was a Vydac 218TP54 RP column (C18, 5 µm, d = 0.46 cm, l = 25 cm) and a flow rate of 1 mL min^-1^ was maintained. Mass spectral data (MS) were recorded on a VG Quattro II spectrometer (ESP ionization, cone voltage 70 V, capillary voltage 3.5 kV, source temperature 80 °C). Data collection was performed using Masslynx software. Structural information from the LC/MS spectrum was visualized using ChemSketch (ACD Labs).

### 2.3 Metal bioleaching

The elemental composition of the two materials used in this study, an iron oxide-rich processing residue from Zn production and a Polish laterite ore was determined via a pseudo total acid digest via aqua regia ^45^ (Table 2) after sieving through a 1000 µm mesh sieve and have been previously characterised ^46^. Briefly, the iron oxide mineral residue primarily consisted of gypsum, quartz, calcite, hematite, willemite, jarosite and franklinite, whilst the laterite comprised of lizardite, forsterite, magnesioferrite, quartz and willemseite. All batch leaching experiments were performed in closed polypropylene tubes (Greiner, Germany) at 28 °C in a vertical shaker.

**Table 2.** Elemental composition of the investigated materials, expressed as mg metal per g of material. Concentrations are mean values and standard deviations derived from chemical analysis in triplicate (N = 3).

| Element | Iron oxide mineral residue (mg g<sup>-1</sup>) | Laterite (mg g<sup>-1</sup>) |
| --- | --- | --- |
| Al | 6.03 ± 0.09 | 1.44 ± 0.11 |
| Cd | $0.287 \pm 0.006$ | $< 0.0003$ |
| Co | $0.020 \pm 0.001$ | $0.164 \pm 0.011$ |
| Cr | $0.337 \pm 0.006$ | $0.412 \pm 0.058$ |
| Cu | $2.95 \pm 0.10$ | $0.001 \pm 0.001$ |
| Fe | $125.5 \pm 4.24$ | $67.9 \pm 3.0$ |
| Mn | $2.45 \pm 0.07$ | $1.36 \pm 0.08$ |
| Ni | $0.050 \pm 0.001$ | $10.3 \pm 0.4$ |
| Pb | $12.5 \pm 0.6$ | $< 0.015$ |
| Zn | $43 \pm 2$ | $0.084 \pm 0.002$ |

For the fractional factorial design of experiments (DOE), five parameters were evaluated: sonication of the material prior to leaching (yes/no), pulp density (5%/20%), pH (2/9), particle size fractions (0.2 mm/1 mm) and the presence or absence of microbial biomass, i.e. separation prior to leaching via centrifugation (yes/no). All subsequent experiments were performed without sonication, a pulp density of 5%, no pH buffering and on material sieved with a 0.2 mm steel wire mesh. After 24 h, the pH was measured and the suspensions were filtered using 0.2 μm syringe filters (Chromafil Xtra, Germany). All experiments were performed in triplicates and control experiments were conducted with demineralized H_2_O and uninoculated MSM (pH 7). The negative controls in demineralized H_2_O and MSM had a pH value of either 2 or 9.5. Siderophores produced by the three strains of *Pseudomonas* were harvested at the previously determined maximum siderophore unit (SU) production time point of 48 h and verified for siderophore concentration using the CAS assay, prior to bioleaching. The siderophore content was calculated according to Equation 1 where A_r_ and A_s_ correspond to the absorbance at 630 nm of the reference (sterile growth media) and sample, respectively ^18^.

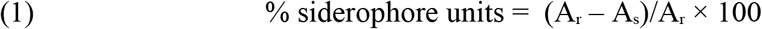

Leaching experiments were also conducted with synthetic siderophore functionalities, using catechol (Sigma Aldrich, Germany), acetohydroxamic acid (Sigma Aldrich, Germany) and a commercial purified siderophore, 1.52 mmol L^-1^ deferoxamine (DFO, Sigma Aldrich, Germany). The functional analogues catechol and acetohydroxamic acid (AHA) were added at both low (0.1 wt/v %) and higher 1 and 10 wt/v % in order to exaggerate differences in leaching and to develop insights into siderophore-metal interactions in these systems.

### 2.4 Chemical analysis

The metal concentrations in the filtrate were in-line diluted with a 1 μg L^− 1^ Rh internal standard and determined by Inductively Coupled Plasma – Optical Emission Spectroscopy (ICP-OES, Varian Vista MPX, US), after appropriate dilution using 1 (v/v)% HNO_3_. Quantification was performed using an external standard series and linearity criteria in the calibration of R^2^ > 0.9990. All reported concentrations exceeded the method detection limit. The pH was measured using a Consort multiparameter analyzer C3020.

## 3. Results

### 3.1 Microbial siderophore production and purification

The siderophore production by *P. fluorescens, P. azotoformans* and *P. putida* was compared under previously reported optimal siderophore production conditions ^18,41^. Similar average growth rates of 0.092 ± 0.009 h^-1^, 0.080 ± 0.001 h^-1^, 0.095 ± 0.008 h^-1^ were observed for *P. putida, P. fluorescens* and *P. azotoformans*, respectively (Figure 1A). Whilst *P. putida* and *P. fluorescens* had similar maximum SU production rates (3.0 ± 0.2 h^-1^ and 2.6 ± 0.8 h^-1^, respectively) compared to *P. azotoformans* (1.6 ± 0.7 h^-1^), near maximal SU units were measured at an earlier time point with *P. putida* (21 h) with respect to the other 2 strains (Figure 1B). No significant enhancement of siderophore production was observed by using a 10 fold higher inoculum concentration (starting OD_600_ 0.2 vs 0.02), highlighting siderophore production was an active process during the growth of these strains. The maximum yield of 75 % SU by *P. putida* in this study is slightly lower than the 83 % and 87 % reported by Sayyed and coworkers ^18^ and higher than the value of 69 % SU for *P. aeruginosa* reported by Shaikh and coworkers ^47^. With its optimal siderophore production, *P. putida* was therefore chosen for further leaching experiments in this study.

**Figure 1.**
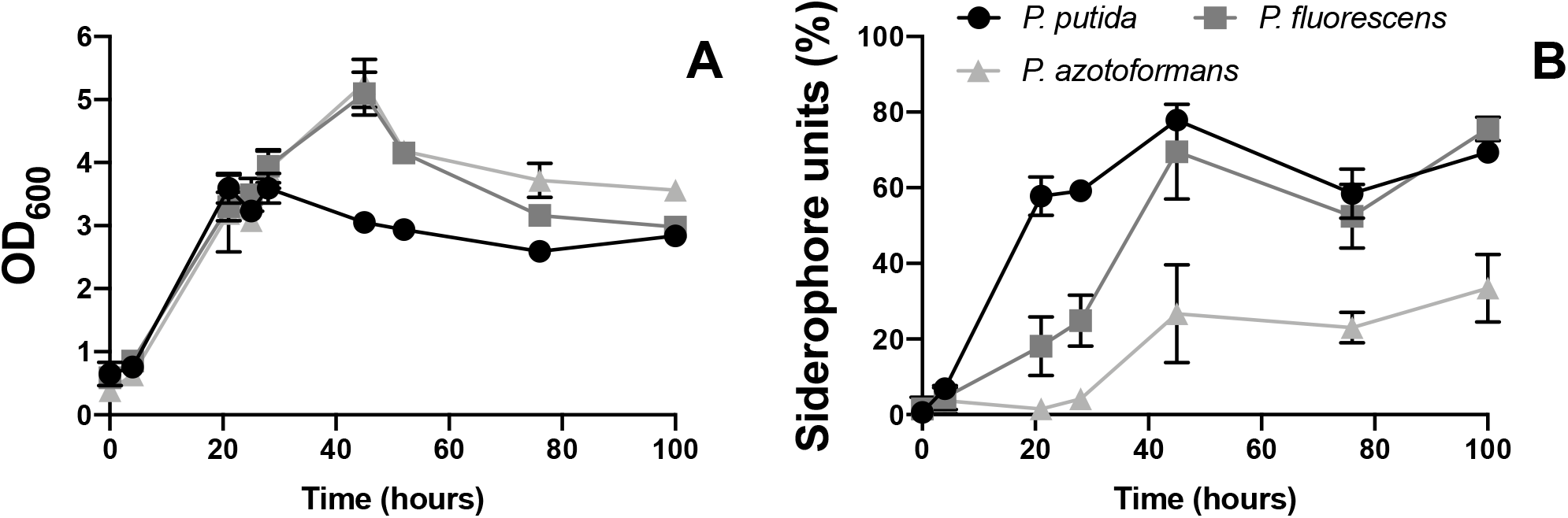
Growth curve (A) and siderophore production (B) by *Pseudomonas putida, Pseudomonas fluorescens* and *Pseudomonas azotoformans* as a function of time. Error bars arise from *n* = 3 independent samples.

To identify the pyoverdine produced by *P. putida* under the experimental conditions of this study, siderophores were harvested from the growth media and analyzed by ESI-MS. Structural analysis of semi-purified siderophores by ESI-MS (Figure S1) showed a predominant molecular mass at *m/z* 1370 and at its double ionization of *m/z* 685, corresponding to a previously identified a pyoverdine-type siderophore, PyoPpC-3B ^48^, with the chromophore group linked to a 9 residues long peptide chain consisting out of Asp-OHbutOHOrn-Dab-Thr-Gly-Ser-Ser-OHAsp-Thr. The abbreviations OHbutOHOrn, Dab and OHAsp represent N^δ^-hydroxybutyryl-N^δ^-hydroxy-Orn, diamino-butanoic acid and threo-β-hydroxy-aspartic acid, respectively. The peptide sequence suggests a metal-binding pocket formed by three moieties; the catecholate of the chromophore, the hydroxamate of N^5^-hydroxy-Orn and β-hydroxybutyric acid, and the α-OH-carboxylate from OHAsp. PyoPpC-3B has been shown to be actively involved in iron acquisition by another closely related strain, *P. putida* C ^49^.

### 3.2 Physiochemical impacts towards metal extraction by biogenic siderophores

To efficiently identify biological (presence of cells) and physiochemical (pH, particle size, pulp density and sonication of the material) parameters that may influence metal extraction from an iron oxide mineral residue by siderophores, a fractional factorial DOE was employed (Figure 2). No metals were extracted in the growth media control, indicating that metals leached were through a combination of siderophore and/ or physiochemical modifications. Whilst sonochemical leaching has been demonstrated to impact (bio)physiochemical parameters and improve (bio)leaching processes, no significant impact on metal extraction was observed in our experiments (*p* = 0.250) ^50^. Aside from copper, a slightly poorer bioleaching performance in the presence of residual *Pseudomonas* cells was observed. Further production of siderophores may have been suppressed due to the initially extracted metals in the pregnant leachate, and the overall metal extraction performance may have been counteracted by their adverse sorption to cell surfaces. Nevertheless, the difference was not significant (*p* = 0.851). The highest metal extracted over all conditions was Zn, with a marked selectivity (900 fold average response Zn vs Fe) over the highest abundant metal in the starting material, Fe (24 wt %). The most significant factor on metal extraction was lowering the pH to 2 (p < 0.05 for Zn), highlighting the importance of proton attack towards metal solubilisation from this material. Lowering the pulp density from 200 g L^-1^ to 50 g L^-1^ generally improved the leaching of all metals analyzed, which is typically reported from previous leaching studies with biogenic acids as it improves reactive processes at the surface interface ^51,52^. With the exception of iron, increasing the total particle size fraction from 0.2 mm to 1 mm hindered leaching, which may represent more facile surface reactions on smaller particle sizes for non Fe metals and has also been reported for inorganic acid leaching ^53^.

**Figure 2.**
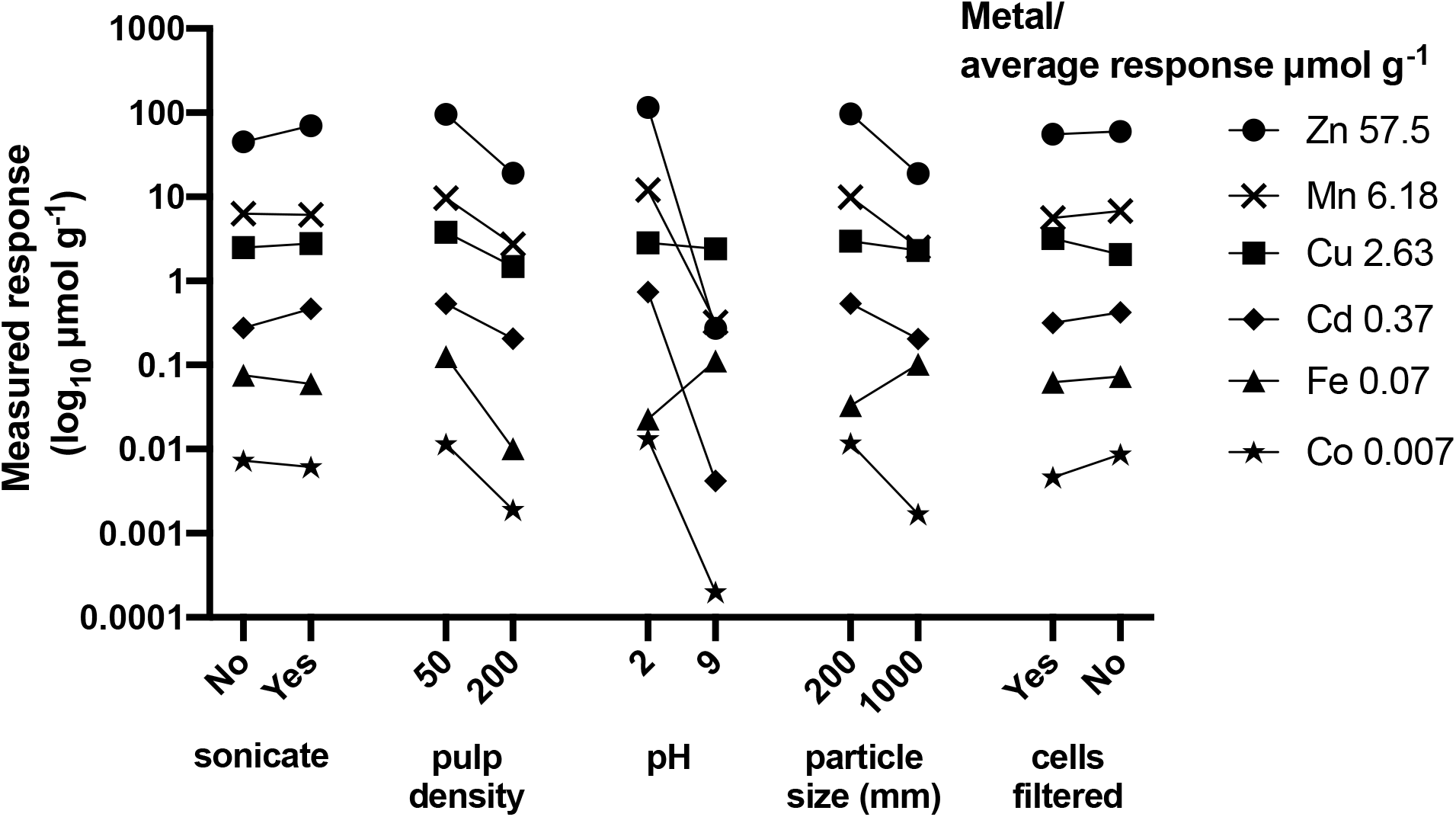
Effect of sonication, solid to liquid (S/L) ratio, pH, sieving mesh size of the substrate and presence of cells on the Zn, Mn, Cu, Cd, Fe and Co extraction from iron oxide mineral residue, presented as its calculated response (log_10_ μmol g^-1^) to each treatment. The average response for each metal is given in the legend

### 3.3 Bioleaching with pyoverdine produced by *Pseudomonas putida*

To further explore pyoverdine-metal extraction mechanisms and to support the initial physiochemical screen with raw siderophore solutions, a series of leaching experiments were conducted with the purified PyoPpC-3B in contact with the iron oxide mineral residue. To augment metal extraction, experiments were sieved with a 200 µm mesh at a pulp density of 50 g L^-1^ without sonication and after removal the biomass of *P. putida* via centrifugation. Whilst dramatically improving the leaching efficiency of Zn, the pH was not changed in order to dissociate the effect of proton activity from pure siderophore activity.

To determine whether pyoverdine purity, concentration or material contact time influenced metal extraction, materials were brought in contact with the lixiviant either directly in the liquid phase after centrifugation of cells, or following a purification step by C18 chromatography and re-dissolving the lyophilized siderophore sample in ultra-pure water at two concentrations over a contact time of seven days (Figure 3A and 3B). The purification step had no effect on the leaching of Mn (p = 0.830) or Zn (p = 0.900) from the iron oxide mineral residue, thus harvested pyoverdines can be applied directly for bioleaching without the need for additional purification steps. Furthermore, expanding the contact time from 1 to 7 days (Figure 3B) only moderately enhanced the leaching of Mn (p = 0.206) and Zn (p = 0.804), showing limited kinetic dependence of bioleaching at neutral pH. The pyoverdine concentration has a certain, although not significant, effect towards Mn (p = 0.173) and Zn (p = 0.346) extraction from iron oxide mineral residue (Figure 3C). Lower concentrations of Al (2.1 ± 0.2 µmol g^-1^) and Cu (1.1 ± 0.1 µmol g^-1^) were also observed after leaching with the higher pyoverdine concentration (Figure 3C). The maximal extraction of Mn (4.6 ± 2.7 µmol g^-1^) and Zn (6.0 ± 2.1 µmol g^-1^) was obtained at the longest contact time of 7 d and highest pyoverdine concentration of 3.6 mM, whereas Cu and Al concentrations dropped at later time points (data not shown), indicative of a reprecipitation event.

**Figure 3.**
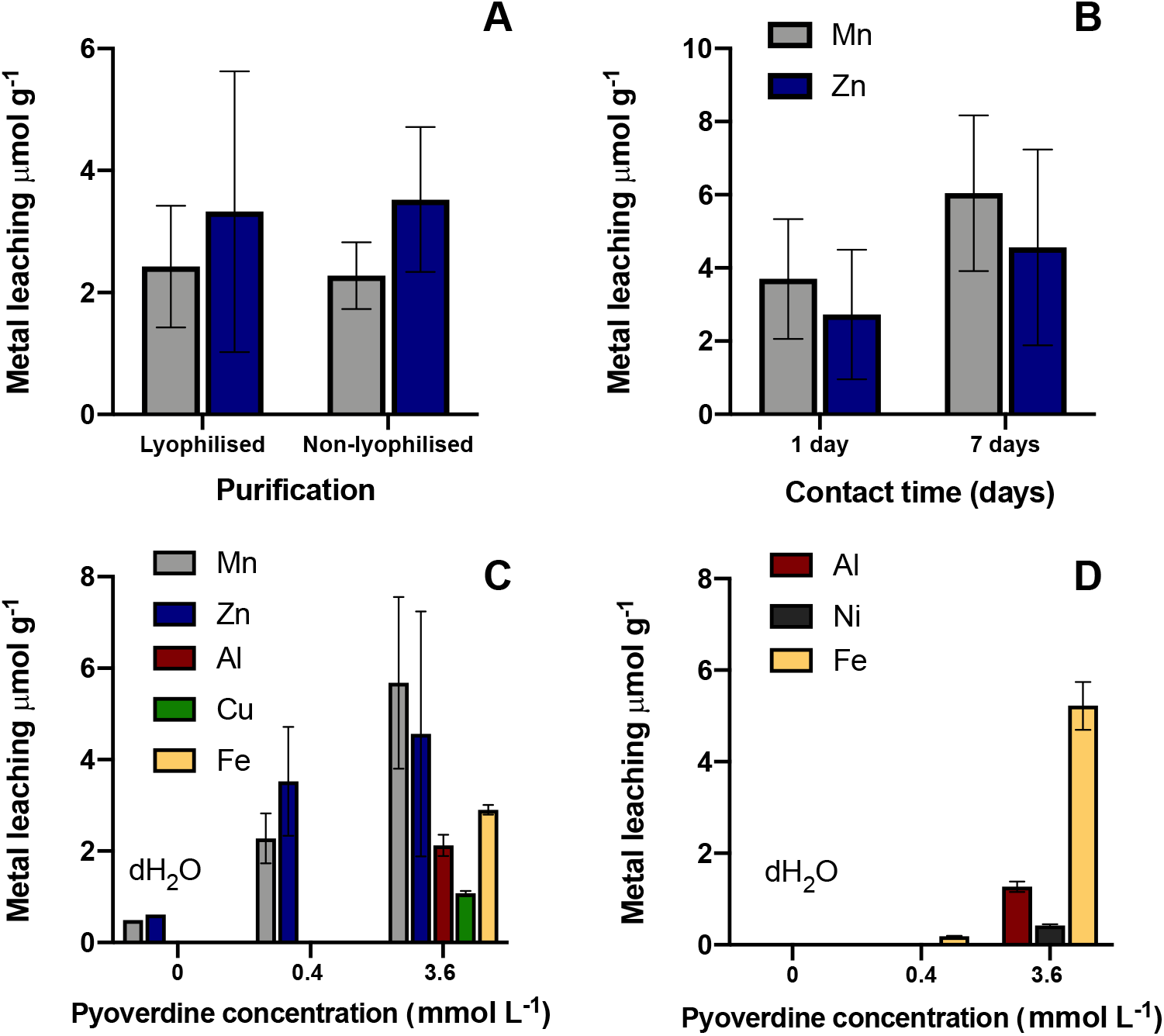
Metal equivalents extracted from low grade mineral residues by PyoPpC-3B. The effect of a purification step to enhance the pyoverdine purity (A), contact time (B) and pyoverdine concentration after a contact time of 1 day with iron oxide mineral residue (C) and laterite (D) on the extracted metal content is shown (*n* = 3).

To explore pyoverdine interactions with other metals and varying mineral phases, leaching experiments with the semi purified pyoverdine were performed on a Polish Ni laterite ore. Similar to the iron oxide mineral residue, higher pyoverdine concentrations favored metal extraction, with 1.3 ± 0.1 µmol g^-1^ Al extracted only at 3.6 mmol L^-1^, coupled to a pH drop from 8.3 to 7.8 (Figure 3D). Ni (0.42 ± 0.03 µmol g^-1^) and Fe (5.2 ± 0.5 µmol g^-1^) were also present after leaching with the higher pyoverdine concentration, consistent with the Ni:Fe ratio of the primary Ni mineral in this material, forsterite. One-way ANOVA with Holm-Šídák post-hoc testing showed an enhancement (p < 0.001) at a pyoverdine concentration of 3.6 mM compared to the control group with only H_2_O. A marked increase of Co extraction was also observed from the laterite using 3.5 mmol L^-1^ pyoverdine (12.5 ± 0.1 nmol g^-1^) compared to 0.4 mmol L^-1^, where Co was below the detection limit (<1 nmol g^-1^). However, these results represent a very low extraction yield and selectivity.

### 3.4 Leaching using synthetic chelating functionalities

In order to gain a mechanistic understanding of the affinity in metal complexation reactions, metal extraction from the iron oxide mineral residue and the Ni laterite ore was evaluated using synthetic chelating functionalities that resemble the chromophore groups found in siderophores. An initial screening was carried out with catechol and acetohydroxamic acid, as representation of catecholate and hydroxamate functional analogues of PyoPpC-3B at 1 wt/v % (133 µmol L^-1^ and 90.8 µmol L^-1^ respectively) ^54^ (Figure 4).

**Figure 4.**
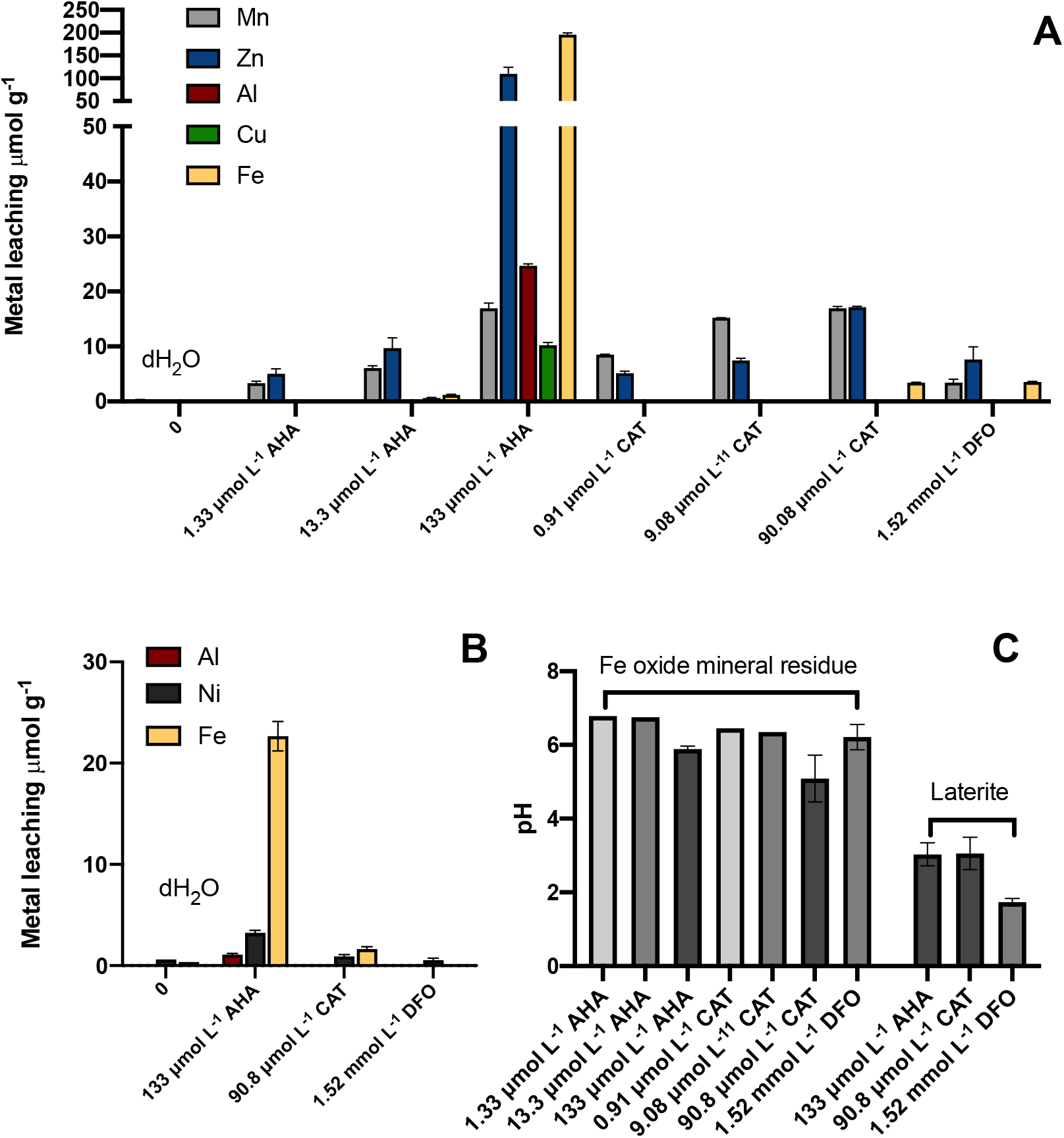
Metal leaching from iron oxide mineral residue (A) and laterite (B) by purified water (negative control), deferoxamine (DFO), acetohydroxamic acid (AHA) and catechol (CAT), with their respective final pH after a contact time of one day. Error bars arise from *n* = 3 independent samples.

In general, as predicted, more pronounced levels of metal extraction were observed using synthetic analogues that were poised at elevated concentrations in comparison to the biogenic pyoverdine. Amongst the different chelating molecules, AHA showed capable of extracting and complexing the largest amount of metals. This was apparent for Zn and Fe in iron oxide mineral residue (Figure 4A) and Fe, Mn and Ni in laterite (Figure 4B). However, the selectivity towards Zn, Mn or Ni over Fe was limited, with relatively high concentrations of Fe being extracted in solution: 170 µmol g^-1^ from iron oxide mineral residue and 23 µmol g^-1^ from laterite. Preferential extraction via acidolysis and subsequent metal complexation by AHA functionalities would suggest that this functional group may have the most pronounced influence towards quantitative bioleaching.

To further explore the impact of hydroxamate and catecholate concentrations on the leaching of metals from the iron oxide mineral residue, leaching experiments were performed at two additional concentrations, 0.1 wt/v % (13.3 µmol L^-1^ and 9.08 µmol L^-1^) and 0.01 wt/v % (1.33 µmol L^-1^ and 0.91 µmol L^-1^) (Figure 4A). Lower catechol concentrations showed a marked improvement of selectivity and Mn extraction. Furthermore, no impact on Mn extraction was observed between two higher concentrations (9.1 and 91 µmol L^-1^), where the biggest difference in the final pH was observed (∼pH5-6.5) (Figure 4C), highlighting that catechol-Mn interactions could not be explained by the pH and could proceed at low stochiometric ratios. With respect to the hydroxamate functionality, a significantly improved selectivity of Zn against Fe was observed at lower hydroxamate concentrations, from a 1:2 Zn:Fe ratio at 133 µmol L^-1^ to 10:1 and 5:0 for 13.3 µmol L^-1^ and 1.33 µmol L^-1^. In this case, the pH shift from ∼pH 5.9-6.8 between the two higher concentrations had a greater role in Zn and Fe extraction (Figure 4A and 4C).

To benchmark PyoPpC-3B against other biogenic siderophores, leaching experiments with a commercial hydroxamate-type siderophore DFO produced by *Streptomyces pilosus* at a concentration of 1 g L^-1^ (1.52 mmol L^-1^) was performed on the iron oxide residue (Figure 4). Whilst Zn leaching was higher for DFO over PyoPpC-3B at 7.65 ± 2,3 µmol g^-1^, a lower selectivity against iron was also observed, with extracted iron at 3.58 ± 0.1 µmol g^-1^. The low mg L^-1^ concentration range of metals leached by PyoPpC-B3 and DFO are comparable with previous reported values from a study using 3 mmol L^-1^ DFO, particularly for Fe (5.2-12.2 µmol g^-1^) that was also present in hydroxide form and at an elevated concentration (7.2-9.2 wt %)^55^.

## 4. Discussion

Bioleaching by heterotrophic microorganisms occurs by one or a combination of the following mechanisms: acidolysis, complexolysis or redoxolysis ^56,57^. Upon contact of the 5 g L^-1^, pH 7.7 PyoPpC-3B lixiviant with the iron oxide-rich mineral residue, the pH of the resulting solution raised from 5.72 ± 0.01 increased to 6.88 ± 0.01 after 1 day. The subsequent pH transition from weak acid to neutral conditions is consistent with the dissolution of Zn^2+^ from the Zn bearing mineral jarosite in the iron oxide mineral residue ^58^ (Figure 3), given according to reaction Equation 2. A similar pH evolution was observed with demineralized water, yet resulted in a lower leaching yield, clearly highlighting that PyoPpC-3B enhanced the metal extraction process. Indeed, results from DOE screening demonstrated that reducing the initial pH (2) through inorganic acid addition could further enhance metal extraction (Figure 2), however no significant difference was observed between the siderophore solution and the acidified siderophore solution, indicating proton attack overrules the siderophore chelating activity.

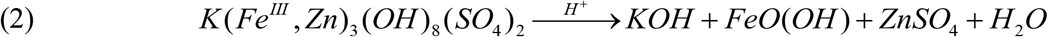

In this study, acidification by acetohydroxamic acid, particularly at higher concentrations highlighted that acidolysis may play a more direct role in metal extraction. Santos and coworkers also showed through pH dependence studies that metal extraction by trihydroxamate is accelerated with increasing acidity of the medium ^59^. Mechanistically, they inferred two parallel pathways: one is a bimolecular process involving the direct attack of the acid on the siderophore analogue complex, the other involves initially the protonation of the trihydroxamate group, followed by a rapid attack of the competing ligand. Conversely, pure complexolysis by siderophores that contain hydroxamate groups is expected to have a higher efficiency at neutral to alkaline conditions. Hence, dissolution and complexation of Al from the laterite in this study is only occurring at neutral pH. The small decrease in pH (from 8.3 to 7.8) during bioleaching can be explained by the release of hydroxonium ions in the pregnant leachate solution (Equation 3).

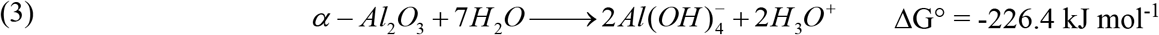

The balance of pH on the added effect of siderophore action is therefore a key parameter to consider towards the application of this technology. Kraemer and coworkers also found that a low pH (< 2) indeed limits metal chelation with siderophores due to a change in the protonation ^55^. Moreover, high H_3_O^+^ concentrations can even cause degradation of the pH-sensitive DFO molecule. Conversely, at pH 2-7, the tertiary amine of hydroxamate moieties in DFO have a lone electron pair, allowing for tris-hydroxamate to coordinate with Fe(III) ^60^. Slightly basic conditions of pH > 8 have also been shown to be preferable for chelation of Pt and Pd by DFO from PGM-rich ores ^55^. In the same way, Neubauer and coworkers showed that DFO could chelate Co^3+^ better than Fe^3+^ at higher pH values ^61^. Whilst preliminary experiments in this study at pH 2 indicated no sign of pyoverdine degradation at early time points and that functional groups were predominantly protonated (data not shown), further work is needed to monitor siderophore stability under these experimental conditions over longer time frames. Identifying whether siderophore degradation (via e.g. ester/amide hydrolysis) or ion exchange with other metals in the pregnant leachate could help to elucidate observations towards Cu and Al reprecipitation from the iron oxide mineral residue over longer leaching time frames.

Redoxolysis can also contribute to the bioleaching process, considering catecholate has a comparable redox potential (ε^0^ = +0.795 V) for e.g. Mn(IV) reduction (ε^0^ = +1.224 V), thus can act as a specific mediator and transform to *o*-benzoquinone ^62^. The dissolution of metal ions from pyrolusite (Equation 4) and manganite (Equation 5) is hereby accompanied by an electron transfer and reduction of Mn(IV) or Mn(III). The relatively high yields of Mn solubilisation in comparison to the lower catecholate concentrations, non-shifted between 16.9 ± 0.34 µmol g^-1^ and 15.2 ± 0.03 µmol g^-1^ for 90.8 and 9.08 µmol L^-1^ respectively (Figure 4), may also give support to an electron shuttling reductive solubilisation mechanism.

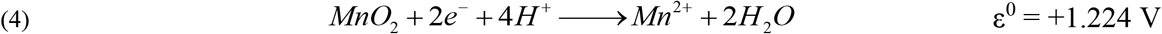

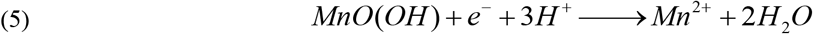

Figure 5 illustrates the central hexadentate metal coordination of the pyoverdine characterised in this study that contains both catecholate and hydroxamate functional groups. A comparison of metal extraction with synthetic functional analogues (Figure 4), pointed towards AHA being the main metal chelating group and that catecholate groups may be involved in the reductive dissolution of Mn oxides in the materials tested in this study. Nevertheless, other non-specific functional groups, as part of the peptide chain attached to C1 of the chromophore group may collectively act as metal cation chelators ^63^, but further work is needed to clarify these observations.

**Figure 5.**
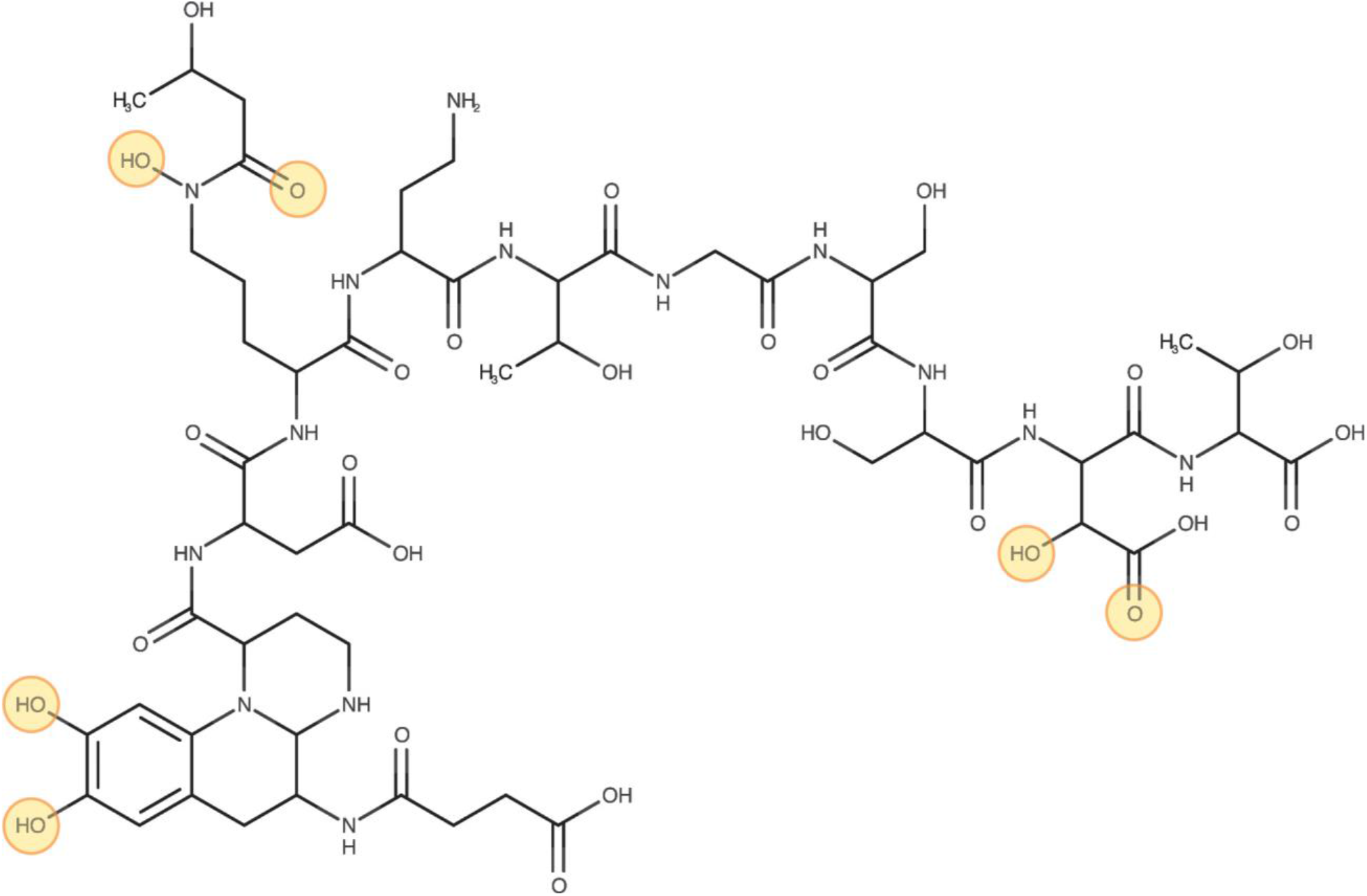
Chemical structure of PyoPpC-3B. The 6 circled functional groups are responsible for metal interaction and together form the metallophore complex

The selectivity in metal complexation is greatly determined by their affinity with functional groups. In mixed metals solutions, competition for siderophore complexation exists between different metal cations, typically Fe^3+^, Al^3+^, Ca^2+^, Cu^2+^ and Zn^2+ 64^. Selectivity of siderophores towards hard metal ions of coordination number 6 became clear from our results (Figure 3), thus in the absence of significant labile Fe^3+^ concentrations, Mn^2+^ and Zn^2+^ may take over its role in complexation with pyoverdine. Both Mn^2+^ and Zn^2+^ possess a similar charge to ionic radius ratio (2: 0.07 nm) that it is determinative for cationic metallophore complexation ^7^. On the contrary, Al(OH)_4-_ follows another geometric complexation as a result of its hydroxyanion speciation.

Pyoverdines are well documented to be selective towards Fe in aqueous mixed metal systems, but little work has looked at the extraction of metals from solid mineral residues. The selectivity of pyoverdines towards Zn and Mn over Fe in this study likely represents a recalcitrance of the Fe mineral phases and/or more labile Zn and Fe for siderophore extraction. Whilst hematite, a principal iron mineral in the iron oxide-rich mineral residue has been previously demonstrated to be susceptible to Fe extraction by siderophores, these studies were performed either through siderophore expression during in situ contact with the material, using an alternate hydroxamate-type siderophore, DFO and/or with nanoparticle fractions ^4^. The larger particle sizes in this study, in combination with its presence within a mixed highly weathered sample may thus impede such processes. This may be compounded by the limited metal binding siderophore substrate. This effect may be less apparent in the laterite system, with lower reported aluminum-hydroxamate formation constants (log K 21.5) ^65^. The higher observed Zn to Fe ratios at lower concentrations of acetohydroxamic acid, where a pH shift from 6 (133 µmol L^-1^) to 6.75 (13.3 and 1.33 µmol L^-1^) occurred may also favor metal complexolysis of Zn over acidolysis, but should be validated with other materials in follow up studies.

## 5. Conclusions

This study demonstrates the selective extraction of Zn^2+^ and Mn^2+^ metal ions, and competitive extraction of Al(OH)_4-_ from a primary and secondary mineral residue, by a hydroxamate and catecholate mixed-type pyoverdine PyoPpC-3B, produced by *Pseudomonas putida* PpF1. We propose the following mode of action of this biogenic macromolecule and its functional analogues: (1) (in)direct metal dissolution via acidolysis (e.g. for Zn from jarosite) (2) reductive mineral dissolution via redoxolysis (e.g. for catecholate groups with Mn oxides) (3) subsequent complexolysis via the chromophore that can further drive metal release from the substrate. Whilst relatively low yields are reported for these materials, metal solubilisation could be further enhanced by increasing siderophore or functional analogue concentrations, and to a lesser extent, the contact time. For further follow up studies, combinations of siderophores and more established bio-hydrometallurgical lixiviants such as biogenic organic acids under weakly acidic conditions could also be of interest to identify synergy between higher leaching yields and selectivity. The separation of the metal-siderophore complex and subsequent metal recovery from the pregnant leachates should also be investigated and recent work has demonstrated that this could be feasible ^66^.

Whilst some work has reported concomitant or alternate metal complexation by siderophores over iron, little work has shown a clear preference for Zn and Mn extraction over Fe. Whilst Al was not selectively leached over Fe in the laterite, the co-leaching of these metals by this particular siderophore has not been reported before and clearly warrants further follow up. Comparison with synthetic functional analogues revealed that AHA has strong metal chelating properties. It is suggested that these hydroxamate moieties in biogenic siderophores also represent the primary route in the metal dissolution and complexation process, although advanced chemical characterization of the metal complexes in the leachate are needed to support this working hypothesis. The stoichiometric excess of Mn extracted against the lower catechol concentrations suggests that reductive dissolution may play an additional role, but further work is also needed to clarify these observations. Finally, the stability of siderophores during the metal extraction process should be evaluated to determine their impact on metal release post extraction and their potential recovery for reuse.

This study has, for the first time, implicated the direct role of siderophores in metal extraction from low grade primary and secondary resources. Whilst currently providing low relative metal yields, it supplies important first steps towards sustainable metal extraction and recovery. Moreover, it provides important insights in siderophore-metal interactions in complex and refractory primary and secondary mineral sources, that also have implications for the acquisition of metals by microorganisms in the environment.

## Acknowledgements

The research was financed by SBO Project SMART (Sustainable Metal Extraction from Tailings) that fits in the SIM (Strategic Initiative Materials) program of Flanders (*HBC*.*2016*.*0456*) and was supported by the EU Horizon 2020 METGROW+ project (Grant Agreement n° 690088) on Metal Recovery from Low Grade Ores and Wastes. We thank Dr. Jasmine Heyse for critically reading the manuscript.

## Author Contributions

AJW conceived the project, designed experiments, assisted in data acquisition, interpreted the data and wrote the manuscript. KF assisted in data acquisition, interpreted the data and wrote the manuscript. YPC performed growth experiments and DOE analysis. SM characterised the pyoverdine structure, assisted in data interpretation and reviewed the manuscript. JV assisted in DOE design and analysis, GDL, NB and TH assisted in project conceptualisation, data interpretation and reviewed the manuscript.

## Notes

### Competing Interest Statement

The authors have declared no competing interest.

