## Supplementary information for "Selective metal extraction by biologically produced siderophores during bioleaching from low-grade primary and secondary mineral resources"

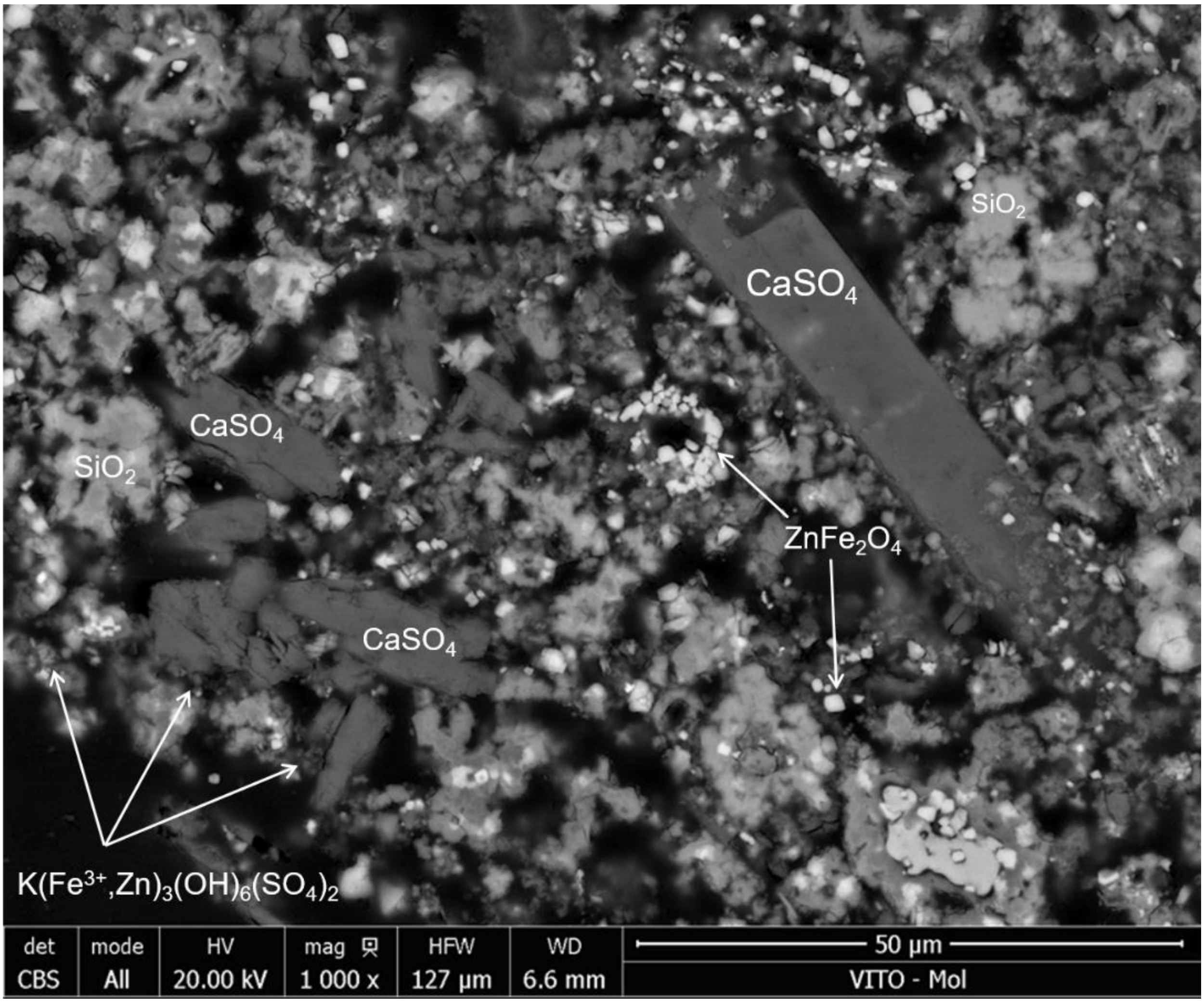


**Figure S1.** Scanning electron microscopy (SEM) image of the iron oxide ore residue. Mineral phases of *gypsum* ($CaSO_4$), *silica* ($SiO_2$), *jarosite* ($K(Fe,Zn)_3(OH)_6(SO_4)_2$) and *franklinite* ($ZnFe_2O_4$) are detected.

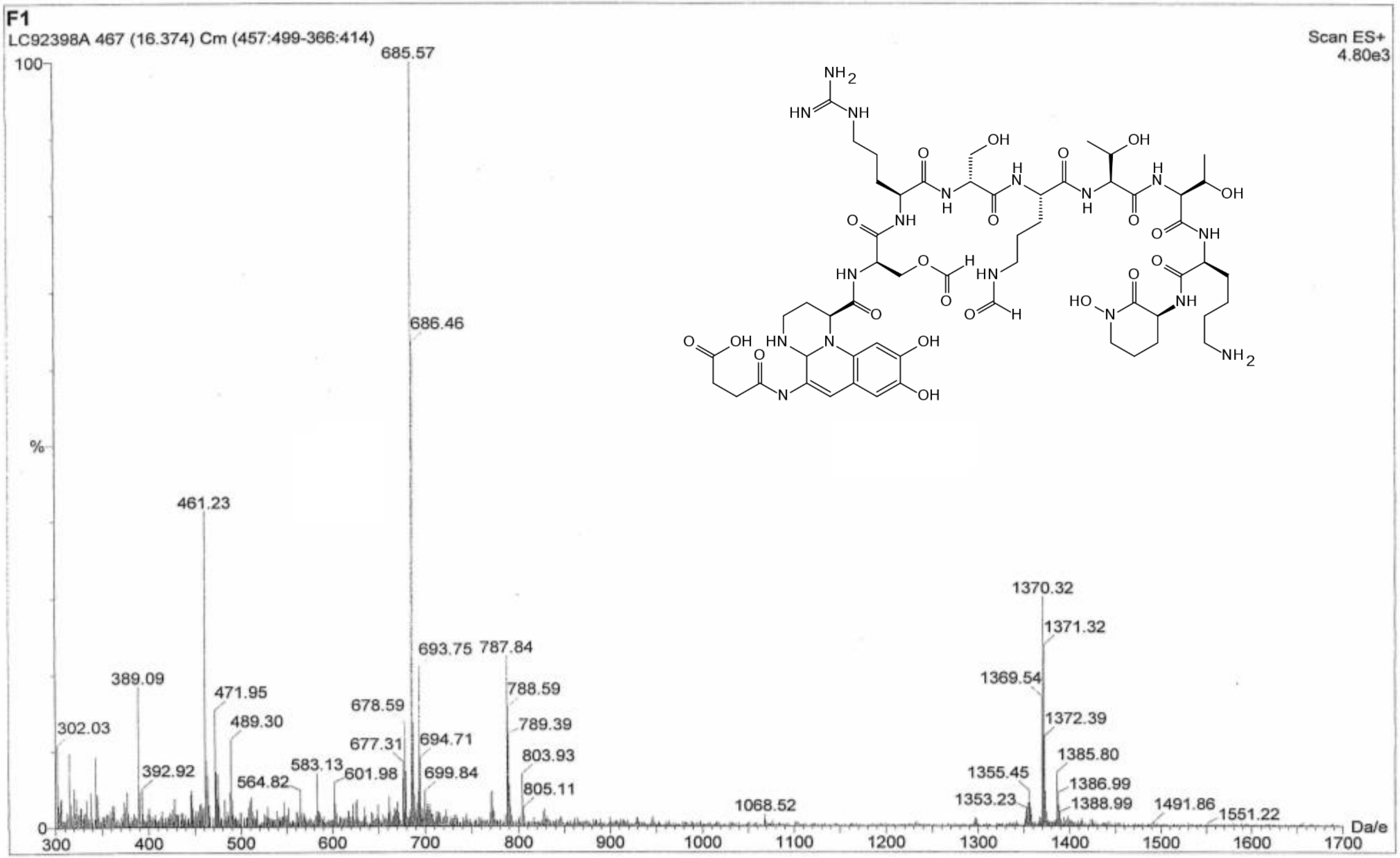


**Figure S2.** ESI-MS spectrum of purified pyoverdine, of which the chemical structure is shown in inset, that was biologically produced from *Pseudomonas putida.*
